# Low HER2 enables dedifferentiation and transformation of normal breast epithelial cells via chromatin opening

**DOI:** 10.1101/2022.09.06.506760

**Authors:** A Hayat, EP Carter, HW King, A Ors, A Doe, SA Teijeiro, S Charrot, S Godinho, P Cutillas, H Mohammed, RP Grose, G Ficz

## Abstract

Overexpression of the human epidermal growth factor 2 (HER2) protein in breast cancer patients is a predictor of poor prognosis and resistance to therapies. Despite significant advances in the development of targeted therapies and improvements in the 5-year survival rate of metastatic HER2-positive breast cancer patients, a better understanding of the disease at an early stage is needed to prevent its progression. Here, we used an inducible breast cancer transformation system that allows investigation of early molecular changes at high temporal resolution. HER2 overexpression to similar levels as those observed in a subtype of HER2 positive breast cancer patients induced transformation of MCF10A cells and resulted in gross morphological changes, increased anchorage-independent growth of cells, and altered transcriptional programme of genes associated with oncogenic transformation. Global phosphoproteomic analysis during the first few hours of HER2 induction predominantly detected an increase in protein phosphorylation. Intriguingly, this correlated with a wave of chromatin opening, as measured by ATAC-seq on acini isolated from 3D cell culture. We observed that HER2 overexpression leads to reprogramming of many distal regulatory regions and promotes reprogramming-associated heterogeneity. We found that a subset of cells acquired a dedifferentiated breast stem-like phenotype, making them likely candidates for malignant transformation. Our data show that this population of cells, which counterintuitively enriches for relatively low HER2 protein abundance and increased chromatin accessibility, possesses transformational drive, resulting in increased anchorage-independent growth *in vitro* compared to cells not displaying a stem-like phenotype. Our data provide a discovery platform for signalling to chromatin pathways in HER2-driven cancers, offering an opportunity for biomarker discovery and identification of novel drug targets.

## Introduction

Metastasis is the main cause of cancer deaths but understanding the root cause of malignant transformation remains poorly understood. Many questions remain unanswered as to what triggers cancer formation beyond DNA mutations in pre-cancerous tissue (Ciccarelli and DeGregori, 2020). Perturbed signalling due to dysregulated phosphorylation of oncogenic proteins is known to alter pathway activity and contributes to cellular transformation (Sever and Brugge, 2015; Hanahan and Weinberg, 2011). Similarly, cell identity and cellular plasticity are phenotypic outcomes of the signalling and epigenetic information in both healthy and disease states (Wainwright and Scaffidi, 2017). Therefore, understanding how an altered signalling environment affects the epigenome and shifts cellular states is crucial in furthering our understanding of cancer formation. Integrating systematic analyses of phosphorylation sites (phosphosites) from global phosphoproteomics data with DNA/RNA sequencing data helps to better understand the functional significance of the signalling effects on chromatin changes. Phenotypic changes that occur during cancer development are driven by changes in the gene expression patterns, which are themselves governed by regulatory states encoded within the nucleoprotein structure of chromatin (Voss and Hager, 2014). The alterations in chromatin structure that lead to differential accessibility to transcription factor binding have been identified as perhaps some of the most relevant genomic characteristics correlated with biological activity at a specific locus (Thurman *et al*., 2012). Nevertheless, the specific regulatory changes driving the transition from normal to transformed cells remain largely unknown.

HER2 positive breast cancer accounts for approximately 20% of all breast cancers (Wang and Xu, 2019). The ability of HER2 positive breast cancer cells to leave the primary tumour site and establish inoperable metastasis is a major cause of death and a serious impediment to successful therapy. Molecular analysis of HER2 positive breast cancer progression is limited by the inability to characterise and catalogue early changes at the onset of transformation. Conventional *in vitro* models (Pradeep *et al*., 2012; Gangadhara *et al*., 2016) can recapitulate the genetics, morphology, therapeutic response and highly transformative nature of the disease. However, they do not allow for the fine tuning and temporal control required to fully assess cellular events leading up to malignant transformation. To overcome this issue, we developed an inducible *in vitro* model of human breast cancer to investigate the mechanisms that drive early transformational changes in HER2 positive breast cancer. The strength of an inducible system lies in that it can recapitulate key transitional states in cancer progression in a controlled manner, permitting isolation of cancer-like cells at defined stages of transformation to catalogue early tumour promoting changes.

Here, we analysed HER2 protein overexpression in a normal diploid, oestrogen, and progesterone negative breast epithelial cell line, MCF10A (Qu *et al*., 2015) to identify global cell signalling and chromatin accessibility changes in the first few hours and days of cellular transformation. In particular, we explored how cell signalling interacts with chromatin to induce transformation as a result of HER2 pathway activation.

### Conditional HER2 overexpression promotes *in vitro* transformation

HER2 overexpression in non-tumourigenic MCF10A cells is a well-established breast cancer model and has been used in numerous *in vitro* studies (Muthuswamy *et al*., 2001; Imbalzano *et al*., 2009). To recapitulate the early transformational events and the stochastic nature of early breast cancer development, we generated a controllable *in vitro* model system by stably transducing a doxycycline-inducible HER2 construct in MCF10A cells (Carter *et al*., 2017). This model allows for the generation of transformed phenotypes in a synchronised and time-controlled manner and is useful for investigating early transformational events using multi-omic analysis (**Fig 1A**). To analyse the range of HER2 expression at the protein level, we cultured cells for 24 hours in five different concentrations of doxycycline, using ranges that have been used previously in inducible expression studies with other proteins (Baron *et al*., 1995; Leitner *et al*., 2014). In our model, a 24-hour induction with 1μg/ml doxycycline resulted in strong HER2 protein expression (**Fig 1B**). When grown in three-dimensional cell cultures, control MCF10A cells (MCF10A^**CTRL**^) formed regular, spherical acini, whereas a majority of MCF10A^**HER2**^ acini were misshapen, with cells budding into the surrounding matrix (**Fig 1C**). HER2 overexpression resulted in significantly increased *in vitro* migratory and invasive potential, as measured by transwell assays (**Fig 1D**) (Xiang and Muthuswamy, 2006; Paszek and Weaver, 2004). Furthermore, MCF10A^**HER2**^ cells displayed a hallmark of *in vitro* transformation, with increased anchorage-independent growth as compared to control cells (**Fig 1E**). Collectively, these results show that HER2 overexpression in MCF10A cells results in phenotypes associated with *in vitro* transformation. Aberrant expression of HER2 is known to induce phenotypes associated with *in vitro* transformation (Seton-Rogers *et al*., 2004) and evokes aggressive tumorgenicity and metastasis *in vivo* (Alajati *et al*., 2013).

**Figure 1.**
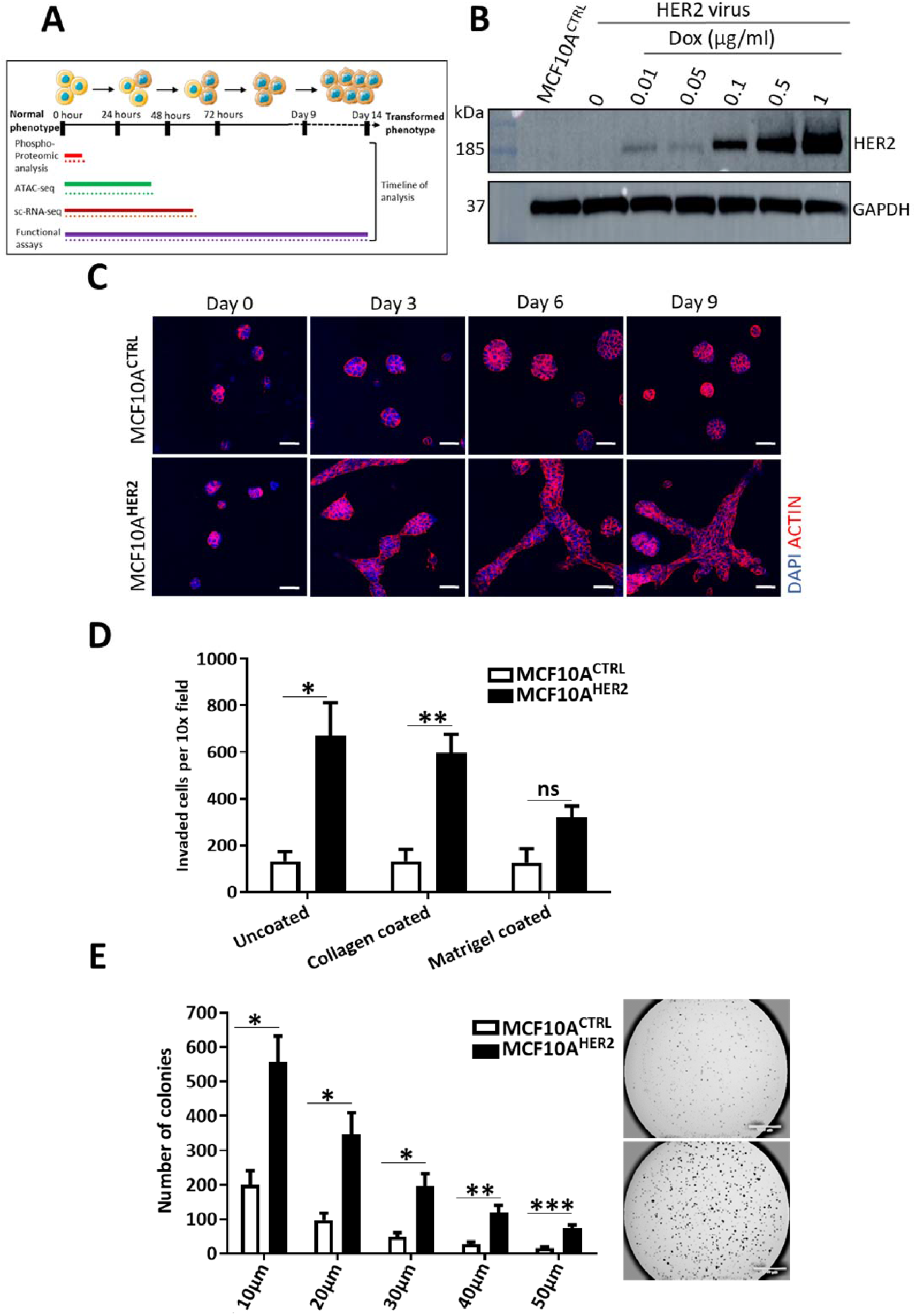
(A) Schematic of multi-omics analysis and soft functional assays performed with their respective timelines as MCF10A cells undergo *in vitro* transformation. (B) HER2 protein expression analysis by western blot in MCF10A cells infected with inducible HER2 lentiviral particles and cultured in various concentration of doxycycline for 24 hours. GAPDH was used as a loading control. N=2. (C) MCF10A^HER2^ and control cells were cultured in 3D cell culture over 9 days. Control cells formed spherical acini which increased in size over time. MCF10A^HER2^ cells formed flat projecting cells of complex masses, typical of transformed cells. Images captured by confocal, LSM 510 microscope. Scare bars represent 50μm. N=3. (D) Cell migration and invasion was analysed through the 8μm pores of transwell membranes over 16-hour period of chemotactic migration towards full serum media. The ability of cell invasion was measured in collagen or matrigel coated transwells. Migration ability was measured in using uncoated wells. Statistical significance was calculated using student’s t-test. Significance is shown as * for p-value < 0.05, ** for p-value < 0.01. N=3. (E) Colony growth of MCF10A^HER2^ and control cells in 0.3% ultra-pure agarose over 3 weeks. ImageJ analysis of 5 different size colonies were quantified. Representative microscopic images of colonies stained with crystal violet after three weeks. Statistical significance was calculated using student’s t-test). Significance is shown as * for p-value < 0.05, ** for p-value < 0.01, *** for p-value < 0.001. Images are at 1.6x magnification. Scale bars represent 1000μm. N=3.

### Phosphoproteomic analysis following HER2 overexpression uncovers signalling changes associated with cancer

HER2 is a tyrosine kinase known to activate a plethora of signalling pathways downstream. To investigate the dynamic changes in the phosphoproteome over time, and the order in which they occur during the phased progression from normal to transformed cells upon HER2 overexpression, we performed an unbiased phosphoproteomic analysis of the early phosphorylation events (at 0.5h, 4h and 7h post HER2 protein induction). The experiment was carried out under standard growth conditions in 2D cell culture, and without serum starving, to be closer to physiological conditions. A GFP-transduced MCF10A cell line was used as a control for doxycycline-only induced changes (MCF10A^**GFP**^). As expected, we observed an increase in HER2 phosphorylation levels in HER2 at T701 phosphosite and its family member EGFR (HER1) at Y1110 phosphosite (**Supplementary Fig 1A).** To filter changes relevant to HER2 induction, we selected only those phosphosites that were significantly changed upon HER2 expression but were not significantly changed in the MCF10A^**GFP**^ cells with a stringent cut-off at log2 fold change for HER2 > 1.5, p-value < 0.05, and log2 fold change for GFP < 5, p-value of > 0.05 (**Fig 2A**). From this refined dataset some potential novel HER2 targets include NUCKS1 (S73) and NUCKS1 (S75), a frequently phosphorylated protein at multiple sites, significantly downregulated at the 4-hour time point (**Fig 2A**) when HER2 protein levels are still quite low as measured by western blotting (**Supplementary Fig 1B**). This protein is known to play a significant role in modulating chromatin conformation (Parplys *et al*., 2015; Grundt *et al*., 2004), and regulates events such as replication, transcription, and chromatin condensation (Ostvold, Anne C., et al, 2001). NUCKS1 phosphorylation at various phosphosites is also known to correlate with breast cancer resistance to retinoic acid, known to have anti-proliferative capacity to several breast cancer cell lines (Carrier *et al*., 2016). Other novel candidates include DDX21, with multiple phosphorylation serine sites at (S164, S168, and S171), which were also significantly enriched in our phosphoproteomic analysis (**Fig 2A**). Since we aimed at investigating the link between signalling and chromatin, we observed that DDX21-bound promoters on average have increased enrichment of active chromatin marks (H3K4me3, H3K27ac, and H39Kac) but are depleted for repressive marks (H3K27me3 and H3K9me3) and promoter-distal (H3K4me1) marks (Calo *et al*., 2015). Some highly phosphorylated phosphosites, which have not been shown to be associated with HER2 protein expression include homeodomain-interacting protein kinase 1 (HIPK1), which is highly expressed in invasive breast cancers (Park *et al*., 2012). SHC1(S246), TTC7A(S182), CDC42EP3(S89), and RIPOR1(S351) were also significantly and stably activated in all the time-points screened, suggesting they may have important roles in the biology of HER2 expressing breast cancer cells (**Fig 2A**). The effect of HER2 overexpression on all proteins was also quantified (**Fig 2B**). Interestingly, of those changes, the 4h time-point showed the largest changes in phosphorylation when HER2 levels are still quite low. Although HER2 protein expression is still low, some of these downstream changes might be present at later timepoints as part of the evolution process.

**Figure 2.**
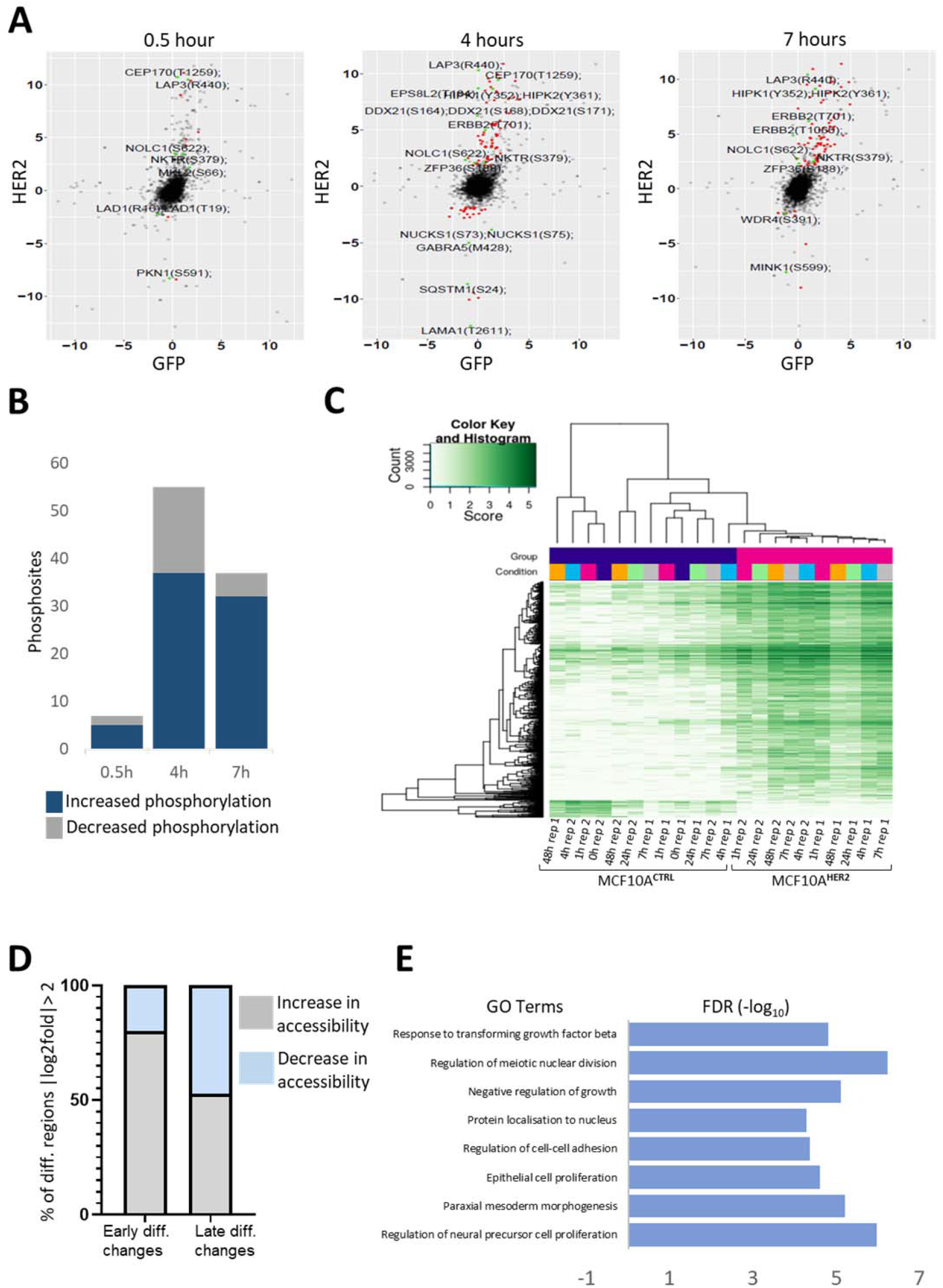
(A) Volcano plots depicting the phosphoproteome upon HER2 protein expression at 0.5-hour, 4 hours, and 7-hour time-points compared to control cells. The statistical significance is shown as (log2 fold change for HER2 > 1.5, p-value < 0.05, and log2 fold change for GFP < 5, p value of > 0.05). The plot contains those phosphosites that are significantly changing upon HER2 protein induction but not significantly changing in the GFP cells at the same time. Those with the highest increase or decrease in fold change are labelled. N=3. (B) Bar graph depicting the number of detected phosphosites increasing or decreasing in phosphorylation in the phosphoproteomic analysis at the indicated time-points analysed. The statistical significance is shown as (log2 fold change for HER2 > 1.5, p-value < 0.05, and log2 fold change for GFP < 5, p value of > 0.05). (C) Differential accessibility (log2 fold change > 0.5, FDR corrected p-value of < 0.05) between MCF10A^HER2^ and control cells, plotted against the mean reads per region. Cells were grown in 3D cell culture from 0-48 hours and ATAC-seq performed on their acini. Heatmap shows chromatin accessibility across all time points for each replicate in cells expressing HER2 or GFP (controls). N=3. (D) Fraction of total regions that are differentially accessible (up peaks) or inaccessible (down peaks) in early or late type comparisons. “Early” time-points represents 0h, 1h, 4h, and 7h data combined. “Late” time point represents 24h and 48h time-points combined. Log2fold > 2, FDR corrected p-value < 0.05. (E) GO categories for biological process for differential peaks that are significantly up ((log2fold change > 0.5, FDR corrected p-value < 0.05) for the Early MCF10A^HER2^ / Early MCF10A^CTRL^ cells.

The low levels of HER2 activation at early time points may closely mimic, at least partially, the early signalling changes occurring in HER2 positive breast cancer patients. The signalling changes of low level HER2 induction has not been performed to date. We re-analysed this data by decreasing the significance threshold to log2fold change > 0.5, FDR corrected p-value of < 0.05 for HER2 expression, but not significantly changing for GFP (Phospho_supplementary_data). This analysis revealed significant changes in phosphorylation in 1045 phosphopeptides over all timepoints in MCF10A^**HER2**^ cells, where the number of phosphosites increased in a time-dependent manner (**Supplementary Fig 1C**).

Using the DAVID Functional Annotation Tool (Huang da, Sherman and Lempicki, 2009), and filtering for all significant changes (log2 fold change > 0.5, FDR corrected p-value of < 0.05) in all the time-points analysed, we identified that mitogen-activated protein kinase (MAPK) signalling pathway to be one of the most enriched cascades in our system (**Supplementary Fig 1D**). The idea that signalling has direct effects on chromatin has already been known, whereby receptor tyrosine kinases can relay extracellular signals by signal transduction pathways to the chromatin (Schreiber and Bernstein, 2002). Signalling pathways, particularly MAPK cascades, elicit modification of chromatin through various transcription factors and chromatin regulators (Clayton and Mahadevan, 2003; Pogna, Clayton and Mahadevan, 2010). Activation of the MAPK pathway ultimately leads to the phosphorylation of transcription factors, which is crucial for gene activation (Treisman, 1996). We hypothesised that the differentially regulated transcription factors and chromatin regulators identified in the phosphoproteomic screen are likely to contribute to chromatin changes mediating the transformed phenotypes. Indeed, our phosphoproteomic analysis revealed significant changes in various transcription factors known to affect chromatin dynamics (**Supplementary Fig 1E**). These chromatin regulators included SIRT1, SOX13, POU2F1, and multiple residues on POL2RA and NCOR1.

In particular, the phosphorylation of JUN at residue S73 could be reconciled by a model in which phosphorylation of JUN triggers dissociation of histone deacetylases (HDACs) and facilitates the rearrangement of chromatin structure (Wolter *et al*., 2008). Based on these results, we then set out to assess, in an unbiased manner, the effects that signalling changes have on the chromatin organisation.

### Identification of two distinct chromatin accessibility landscapes within HER2 induced transformation

To investigate the interplay between signal transduction pathways and chromatin dynamics, we used an assay of transposase-accessible chromatin using sequencing (ATAC-seq) to determine the genome-wide chromatin accessibility landscape in the acini of MCF10A cells in a time-dependent manner (0-48 hours) by isolating cells from 3D cell culture. Principal component analysis (PCA) separated the samples into two groups, “early” (0h, 1h, 4h, and 7h time-points) and “late” (24h and 48h time-points) (**Supplementary Fig 1F**). We selected these conditions with the aim to encompass time-points relevant to both types of analysis. The 0h, 4h and 7h time-points were chosen to characterise early chromatin changes triggered by signalling. The late conditions were selected to detect the resulting delayed chromatin changes occurring later in the process of transformation. We identified 17,868 significant changes between MCF10A^**HER2**^ cells relative to control cells (T0 starting population before HER2 protein induction) over the time course, which showed an increase in accessibility in MCF10A^**HER2**^ cells relative to controls (**Fig 2C & supplementary Fig 2A**). We assessed differential accessibility between early and late groups and observed that a much larger fraction of regions, with > 2-fold difference relative to T0, were enriched in the early group compared to in the late group (75% vs 44%, respectively, **Fig 2D**). Conversely, only ~2.9% of peaks were >4-fold more accessible in the early group and ~6.5% in the late group, which we define as “hyper-accessible” chromatin states (**Supplementary Fig 2A**). Even though the numbers of hyper-accessible versus hypo-accessible regions (which lose accessibility > 4-fold) did not show a stark difference, the overall number of accessible regions following HER2 expression outnumbered inaccessible regions. This shows that there is an increase in chromatin accessibility during the early stages of transformation (**Fig 2D**). Therefore, this might suggest that the first adaptive response to oncogenic HER2 signalling is altered chromatin accessibility to induce differential gene expression. Subsequently, the changes in chromatin accessibility even out in the later time points, with the number of hypo-accessible regions even exceeding the hyper-accessible ones at late time points, which could indicate that cells have reached an equilibrium. (**Supplementary Fig 2A**).

Next, we performed functional enrichment analyses [Gene Ontology (GO) terms] for upregulated peaks in the early HER2 signature (**Fig 2E**). The regions with increased chromatin accessibility at all times analysed were enriched for GO terms associated with response to transforming growth factor, cell-cell adhesion, epithelial cell proliferation, morphogenesis, and regulation of neural precursor cells. The differentially accessible regions upstream of the transcriptional start site (TSS) were largely gene distal, with relatively few promoter-proximal regions (**Fig 3A**). To probe how the observed changes in cell signalling can underlie transcriptional and/or epigenetic control during cellular transformation, we examined transcription factor binding motifs that were significantly enriched in relation to all differential ATAC-seq peaks. The most significantly enriched motifs in the accessible chromatin regions as a result of perturbed HER2 expression were CEBP, HLF, ATF4, and CHOP (**Supplementary Fig 2B**). We also observed significant enrichment of motifs for all the time-points analysed for inaccessible peaks corresponding to closed regions, which included ATF3, AP-1, BATF, FRA1, JUNB, FRA2, and NFkB (**Fig 3B**). Previously it has been shown that enrichment of AP-1 family member motifs is associated with increased accessibility (Hardy *et al*., 2016). There was some overlap between the family members of transcription factors identified in the phosphoproteomic screen and ATAC-seq motif analysis including NFkB, JUN, ATF1, JUND, and AP-1 (**Supplementary Fig 2C**). The transcription factors found in our motif analysis associated with accessible chromatin are known to be involved in several cancer types including breast, lung, endometrial and prostate cancers with a more aggressive phenotype (Detry *et al*., 2008).

**Figure 3.**
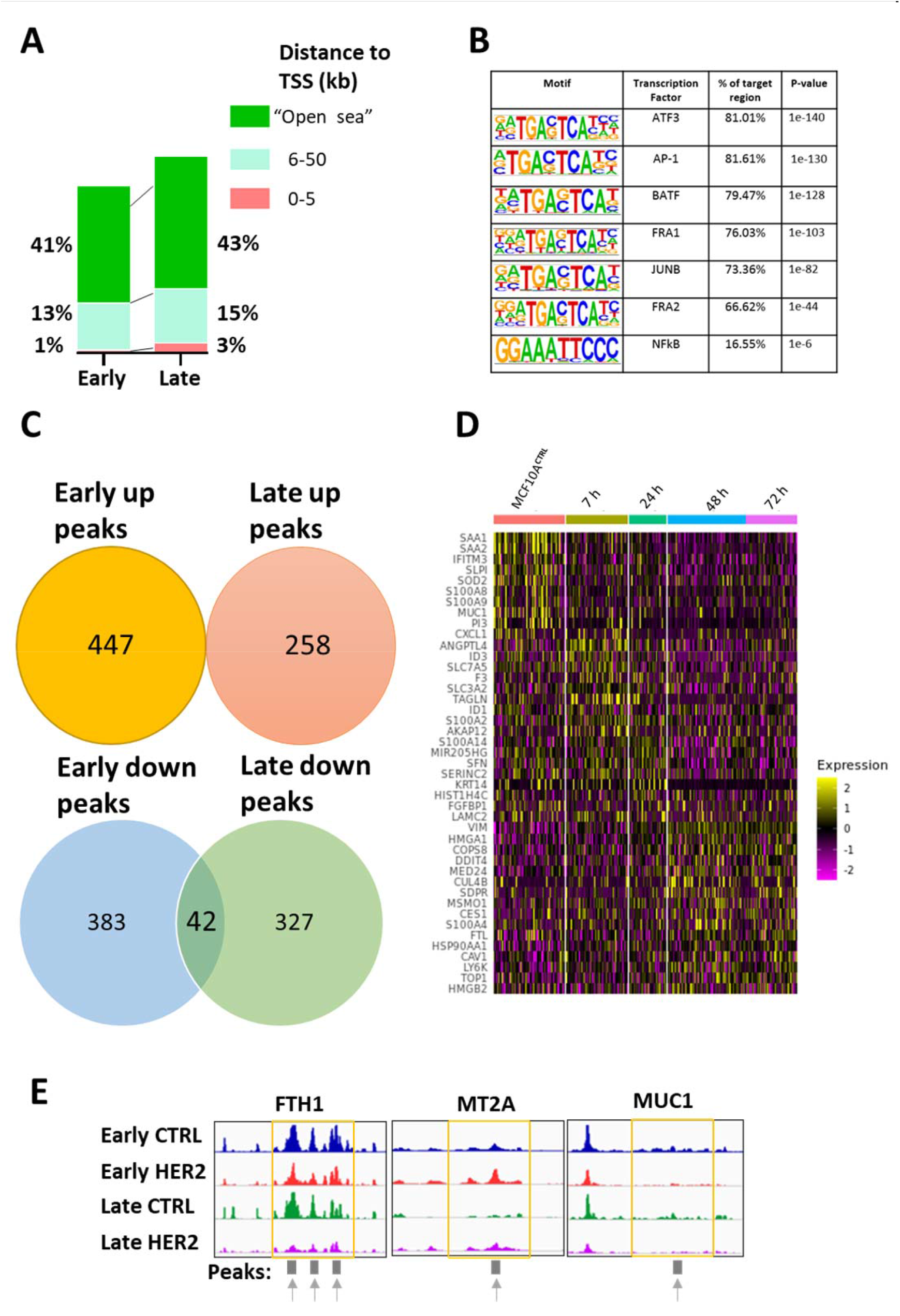
(A) Distance to closest transcriptional start sites (TSSs) of all differentially accessible regions in the early and late cell types. The graphs represents only those regions that are upstream of the TSS. “Open sea” refers to regions that are at least 50 kb or more upstream of the TSS. (B) Enrichment of transcription factor recognition sequences in differential ATAC-seq peaks comparing MCF10A^**HER2**^ and control cells based on HOMER analysis. Down peaks = log2fold < −2, FDR corrected p-value value < 0.05. (C) Venn diagram showing the number of differentially accessible regions that are shared between the up (open) and down (closed) peaks in the early and late samples. Up peaks = log2fold >2, FDR corrected p-value < 0.05. Down peaks = log2fold < −2, FDR corrected p-value value < 0.05. (D) Single cell RNA sequencing was performed in 2D cell culture on MCF10A cells with HER2 induction from 0 to 72 hours (3 days). Heatmap summarises some of the most highly and lowly expressed genes with the induction of HER2 gene. (E) Insertion tracks of samples at example regions. This signal is an average signal of 3 replicates of combined time-points into either “early” samples or “late” samples. Differentially open regions are marked with arrows.

We next examined whether peaks were shared between those that were opening (more accessible) and those that were closing (less accessible) between the early and late groups. We found that there was a small overlap between early and late inaccessible peaks but none between the accessible peaks (**Fig 3C**). This suggests that increasing accessibility is dynamic during transformation, and that sites with early loss in accessibility relative to T0 could potentially have driving roles in the population drift. We further examined the genomic distribution of the differentially inaccessible chromatin of the overlapping regions, which showed most genomic regions were associated with two nearby genes (**Supplementary Fig 2D**). Namely, some of the common differential regions correlated with genomic location of FBN2, whose genomic chromosomal coordinates were found to be matching with the promoter region of the FBN2 gene. This gene was found to have aberrant promoter methylation in a number of cancers (Hibi *et al*., 2012) (**Supplementary Fig 2E).** Other regions included RIMS2, known to be associated with particularly aggressive breast cancers (Zhang, L., Liu and Zhu, 2021) and APIP, which binds HER3 receptor, leading to the heterodimerisation between HER2-HER3 and resulting in sustained activation of downstream signalling (Hong *et al*., 2016). No differentially accessible region was found to be promoter proximal, as all the regions were at least 5 kb upstream of the transcriptional start site (TSS) (**Supplementary Fig 2F**).

To elucidate the heterogeneity in gene expression between subpopulations of cells in light of the pervasive chromatin opening we identified, we performed single-cell RNA-seq following induction of HER2 overexpression over 72 hours. Cells were grouped according to their time-point by UMAP dimensional reduction. Although there is no distinct separation between the time-points, there is a trend in clustering of MCF10A^**CTRL**^ versus HER2 expressing cells (**Supplementary Fig 3A**). Seurat clustering found differentially expressed features and separated them into four groups, with cluster 0 enriching in the MCF10A^**CTRL**^ population, wand cluster 1 associating with the highest HER2 expression (**Supplementary Fig 3B)**. As expected, we observed a time-dependent increase in HER2 gene expression (**Supplementary Fig 3C**). There is a consensus that high HER2 expression is associated with stem-like phenotype (Oliveras-Ferraros *et al*., 2010), however, much controversy remains on whether stemness and high-grade tumours are highly correlated with each other. Some studies have suggested a strong correlation between stemness and high oncogene expression, while others reveal little relationship (Poli et al., 2018; Simeckova et al., 2019). We identified clear transcriptional signatures of oncogenes associated with breast cancer progression such as the time-dependent increase of *ALDOA* (a gene that increases *in vitro* spheroid formation and increases abundance of cancer stem cells) (**Fig 3D and Supplementary Fig 3D**); *LAMB3*, which mediates invasive and proliferative behaviours by PI3K-AKT signalling pathway (Zhang, H. *et al*., 2019), as well as the decrease of genes like *MUC1* conversely upon HER2 overexpression, whose downregulation is linked to stem-like phenotype (Stingl, 2009a). While the expression of *ID3* is also associated with stemness (Huang *et al*., 2019) this pattern was not found in our data, suggesting that these processes overlap only partially.

Genome browser tracks of early and late HER2 samples show the relative accessibility of some regions associated with the indicated gene, with arrow marked regions indicating differentially open regions (**Fig 3E**). Ferritin heavy chain *(FTH1)* gene, which displays sharp decline upon HER2 expression (**Fig 3E & supplementary Fig 3D**) is also associated with inaccessible chromatin, as shown by the scRNA-seq and ATAC-seq datasets. Low *FTH1* expression is known to make breast cancer cells radiosensitive, and its higher expression is correlated with radioresistance (Tirinato *et al*., 2021). An in-depth analysis of *FTH1* expression in HER2 positive clinical samples may improve the efficacy of radiation treatments.

### Sustained low HER2 expression facilitates dedifferentiation and confers stem-like traits

MCF10A^**HER2**^ cells exhibited heterogeneous capacity for anchorage-independent growth when measured by their ability to form colonies in semi-solid medium, in that a significant proportion of MCF10A^**HER2**^ cells were able to form cell aggregates, with a > 2-fold increase in colony forming units compared to control cells (**Fig 1E**). We hypothesised that cells possessing the ability to form colonies under anchorage-independent growth conditions are a selection of aggressive cells out of the total number of cells seeded. Conversely, the proliferative but non-malignant cells that often dominate any heterogeneous parental cell line would be selected against under these conditions. We evaluated whether anchorage-independent growth correlated with reprogramming-associated heterogeneity by testing the expression of proteins found in mammary epithelial stem cell hierarchy by flow cytometry (Stingl, 2009b), in which it has been shown that breast stem cells are characterised by MUC1^**-ve**^, EpCAM^**low**^, and CD24^**low**^ expression (**Fig 4A**). We therefore evaluated whether HER2 overexpression could enrich for cells with functional stem-like properties based on these three markers and found that this stem-like phenotype is enriched in MCF10A^**HER2**^ cells, as a large proportion of cells lost the expression of MUC1, EpCAM, and CD24 (**Fig 4B, Supplementary Fig 3E, and supplementary Fig 4**). Since our population is heterogeneous due to differing number of copies of the lentiviral HER2 construct, and we have the same amount of doxycycline used to induce the oncogene, the upper threshold of expression of HER2 will depend on the transgene copy number. We therefore hypothesised that stem-like markers would be positively correlated with HER2 levels in our heterogeneous population, i.e., cells having many HER2 copies would also be more likely to express stem-like markers. Surprisingly, we found that cells expressing relatively low HER2 levels had the most pronounced stem-like phenotype compared to other flow sorted populations of cells with increasing levels of HER2 (**Fig 4B**). We confirmed the different levels of HER2 protein expression after sorting cells into three compartments of low, medium, and high HER2 expression by western blotting, which correlated as expected (**Fig 4C**). Next, to determine the transformational potential of these cell types by measuring anchorage-independent growth, we flow sorted MCF10A^**HER2**^ cells into the three different cell populations and paradoxically found that low HER2 expressing cells had increased transformational potential relative to the other populations of sorted cells (**Fig 4D**). We thought that high HER2 cells may be undergoing oncogene-induced senescence (OIS), thus resulting in reduced colony formation compared to other cell types. To confirm this, we measured proteins implicated in OIS but found no significant increase in OIS markers in the high HER2 cells compared to other populations, indicating other biological effects being responsible for the lower capacity in anchorage-independent growth of high HER2 expressing cells (**Supplementary Fig 3F**). It is possible that high oncogene expression induces cancer cells to dormancy that is associated with loss of ability to self-replicate and differentiate (Bellovin, Das and Felsher, 2013).

**Figure 4.**
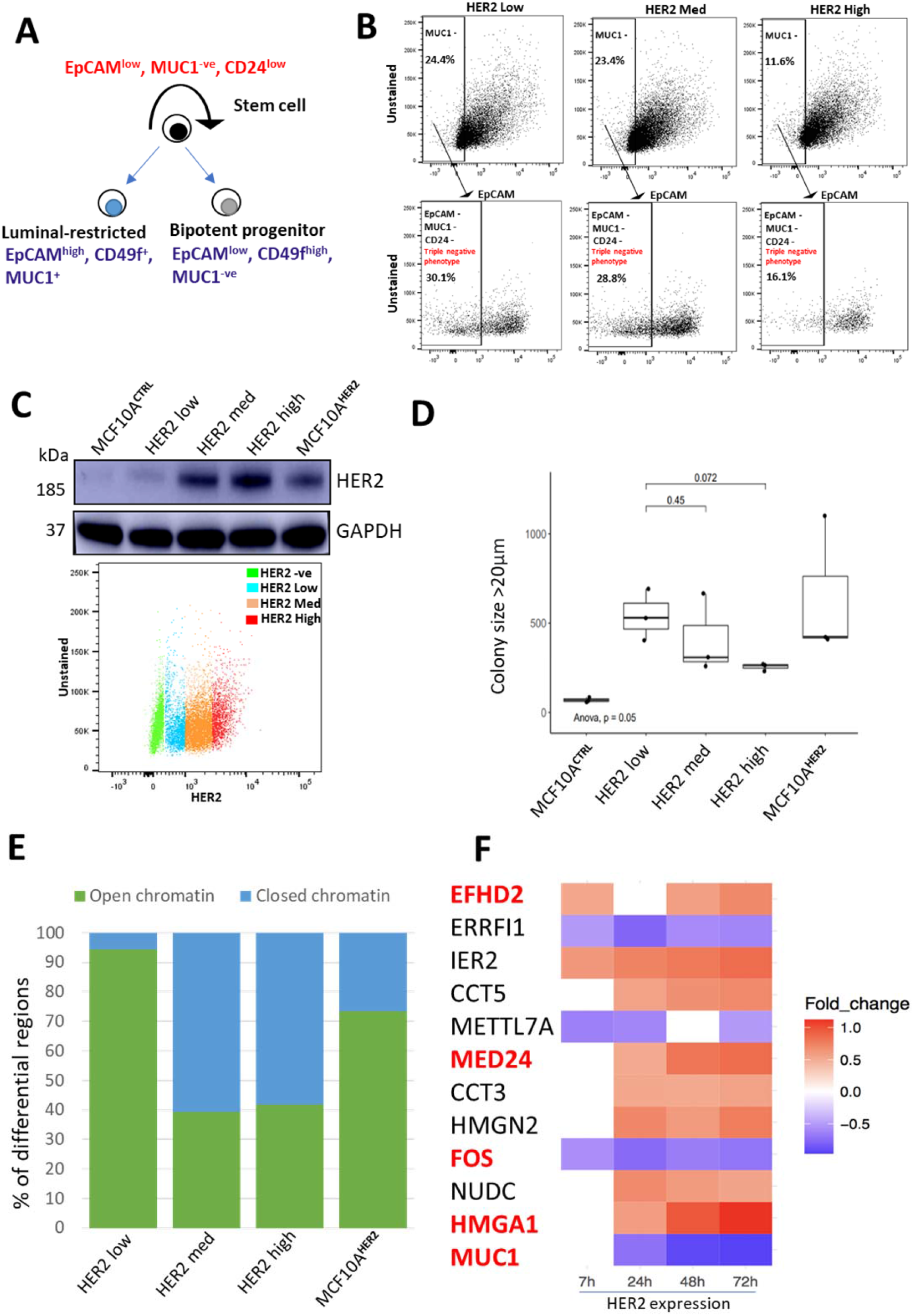
(A) Proposed simplified breast epithelial hierarchy present in human mammary glands. (B) Cells were analysed by flow cytometry and HER2 positive cells were separated into three subpopulations of low, medium, and high HER2 overexpression as indicated. The enrichment of stem markers is shown as a proportion of the total number of cells exhibiting MUC1 ^-ve^ and EpCAM ^-ve^ phenotype. The proportion of cells shown here show the overlap between MUC1 -ve and EpCAM -ve cells, all of which were subsequently 100% CD24 -ve. N=3. (C) MCF10A^**HER2**^ cells were flow sorted into the labelled subtypes and HER2 expression analysis by western blot in MCF10A cells. GAPDH was used as a loading control. The bottom 20% of HER2 expressing cells were labelled as “HER2 low” cells (blue), the top 20% of HER2 expressing cells were labelled as “HER2 high” cells (red). The middle population of 35% were labelled as “HER2 med” (orange). HER2 negative cells are highlighted in green based on HER2 negative control cells. N=3. (D) HER2 expression was induced for 3 days, and cells were sorted based on HER2 expression into low, medium, and high HER2 expression. 5000 cells from each condition were plated into ultra-pure agarose to investigate there *in vitro* transformative potential. Results are plotted as box plots from three biological replicates. Student t-test was performed to compare “HER2 med” and “HER2 high” groups to the “HER2 low” group, p-values are displayed on the graph. One-way anova test was performed to show statistical significance. N=3. (E) MCF10A^HER2^ were sorted into the three subtypes. ATAC-seq libraries were prepared and sequenced. DiffBind was used to analyse the differentially accessible regions and plotted as percentage of open or close regions. N=3. (F) Heatmap shows genes of interest that are consistently differentially expressed in at least 3 of the 4 time-points analysed upon HER2 overexpression. Blue rectangles show genes that are downregulated, red rectangles represent genes that are upregulated. The white rectangles show lack of differential expression for that specific time-point. Only those genes are listed here if the statistical significance had an FDR corrected p-value of < 0.05. Importance of genes highlighted in red are mentioned in the text.

Since we found that chromatin opening was the feature associated with early signalling to chromatin response, we wanted to know if this was reflected in the phenotypic heterogeneity, in particular low versus high HER2 levels. To this end, we used ATAC-seq to determine the genome-wide chromatin accessibility landscape in the five different populations of cells (MCF10A^**CTRL**^, low HER2, medium HER2, high HER2 and MCF10A^**HER2**^ cells). We analysed these data by comparing each cell type to the control cells (MCF10A^**CTRL**^) and comparing the percentage of differentially accessible regions between the cell types. We found that low HER2 expressing cells exhibited the highest percentage of chromatin opening compared to other cell populations (**Fig 4E**), confirming that the phenotypes associated with invasiveness and anchorage independent growth were driven by molecular features in stem-like cells and opening of chromatin. To put the magnitude of these chromatin differences in context, i.e., the differential accessibility between low HER2 and high HER2 expressing cells, we found that a dramatic ~95% of peaks were accessible in low HER2 cells. Conversely, only ~42% of the peaks were open (accessible) in the high HER2 cells. Overall, these data indicate that a sharp increase in HER2, which may result in triggering cell intrinsic defensive systems, whereas a low-level sustained presence of HER2 can shift cell identity, via chromatin remodelling, towards tumour-promoting phenotypes.

We found that a subset of these scRNAseq-unique differentially expressed genes that were either upregulated or downregulated in multiple time-points were also associated with heterogeneity of breast cancer, related to cancer progression and stem cells (**Fig 4F**). For example, expression of *HMGA1*, which is known to promote breast cancer angiogenesis through the transcriptional activity of *FOXM1* (Zanin *et al*., 2019), increased in a time dependent manner (**Fig 4F**). On the other hand, expression of *FOS*, a pro-proliferative transcription factor, which has been validated in breast tumour samples and is highly expressed in relapse samples and treatment failures (Vendrell *et al*., 2008) was found to be downregulated at all time points (**Fig 4F**). Intriguingly, high proliferation rates as a result of *FOS* expression can lead to improved outcomes for patients with breast cancer, as it could lead to higher apoptosis-effector gene expression (Fisler *et al*., 2018). Our data also show the time-dependent increase of *EFHD2*, a gene linked with EMT transition and metastasis (Fan *et al*., 2017). *MED24*, a subunit for the mediator complex of RNA polymerase II, is known to be a downstream target of HER2 and may be a critical gene required for cancer development (Liu *et al*., 2019).

## Discussion

In this study we addressed the question of what the earliest molecular changes are at the interface between increasing oncogenic HER2 signalling and chromatin accessibility in a non-transformed breast epithelial cell line. Overexpression of the HER2 oncogene in breast epithelial cells resulted in some unexpected changes in cellular phenotypes. Namely, we observed an inverse relationship between HER2 levels and tumourigenic properties *in vitro*, where cells expressing a sub-threshold amount of HER2 protein exhibited increased anchorage-independent growth. This was also associated with features of dedifferentiation towards breast stem cell identity. Among the expected features, MCF10A^**HER2**^ cells underwent *in vitro* transformation as evidenced by increased anchorage-independent growth accompanied by the formation of spindle-like conformations in 3D cell culture (**Fig 1B and D**). These findings are concordant with other studies where loss of cell polarity following HER2 overexpression has been described (Ortega-Cava and et al, 2011; Hartman Z., 2012; Xiang and Muthuswamy, 2006b).

We propose that a sub-threshold level of HER2 protein has the ability to elicit activation of signalling pathways that directly impact on chromatin to drive dedifferentiation and survival and to enhance transformation. Although high levels of oncogenic expression are an important biomarker in diagnosing HER2 positive breast cancer, our data support the hypothesis that even low levels of HER2 protein expression can be associated with disease aggressiveness, poor patient outcome and therapeutic resistance (Gilcrease *et al*., 2009). The mechanism of why low HER2 expressing cells can be aggressive and its prognostic value has not been sufficiently evaluated. Our data show that the subset of low HER2 expressing cells likely use changes in chromatin state as their route for cellular transformation (**Fig 4E**); the accessible chromatin induced by low level HER2 signalling may continuously predispose cells to secondary additional hits required for metastasis and therapeutic resistance (Denny *et al*., 2016). The resulting chromatin changes via low HER2 expression may create a lasting and highly transformative state.

Across the different subtypes of breast cancers, and in particular HER2 positive breast cancer, loss of differentiation is associated with lower patient survival and aggressiveness (Margaryan *et al*., 2017; Pupa SM., et al. 2021, 2021). However, in low HER2 expressing cells the correlation between dedifferentiation and aggressiveness remains unclear. Stem marker signatures drive cancer growth, and their inhibition delays it (Rudin *et al*., 2012). Several known stem markers, including the EpCAM, MUC1 and CD44 signatures, promotes transformation and tumour progression (Stingl, 2009). Our data suggest an alternative model in which dedifferentiation is paradoxically driven by low levels of HER2 protein expression and creates a programme of stem cell marker expression that drives transformational ability (**Fig 4B**).

We observed leucine aminopeptidase 3 (LAP3) to be significantly activated in our phosphoproteomic screen in all of the time-points analysed in MCF10A^**HER2**^ compared to MCF10A^**GFP**^ cells (**Fig 2A**). LAP3 is known to play a critical role in breast cancer cells by regulating migration, invasion and is associated with metastasis (Fang *et al*., 2019). In addition, we found that phosphorylation of nucleolar and coiled-body phosphoprotein 1 (NCOL1) at residue S622 was also significantly increased all time-points (**Fig 2A**). This protein is found to be highly expressed in nasopharyngeal carcinomas (NPC) (Hwang *et al*., 2009) and in breast cancer cells (Sacco *et al*., 2016). The consistent and highly stable activation of these two proteins may serve as potential biomarkers for late-stage disease and provide important targets for antimetastatic therapeutic targets. Furthermore, zinc finger protein (ZFP36) is correlated with lower tumour grade breast cancer. Interestingly, we find that ZFP36 (S188) is significantly activated in the 4- and 7-hour time-points but not in the earlier 30-minute time-points (**Fig 2A**), indicating that low HER2 expressing cells prefer a programme of signalling phosphosites associated with worse patient outcome (Canzoneri *et al*., 2020).

The morphological changes in breast cancer models are often used to indicate the high transformational characteristics of those cells (Petsalaki E. et al., 2021). We found that proteins associated with aggressive basal-like phenotype were found to be increased in our phosphoproteomic screen, which included ADGRA2 (S1079) and DENND4C (S1250). This shows that the morphological changes observed in our system (**Fig 1B**) were likely due to HER2 induced transformation.

It is possible that the intrinsic heterogeneity found within the tumour population may be preventing specific patterns from emerging in a bulk RNA-seq analysis. It is known that differential downregulation of IFITM family members is associated with resistance maintenance following anti-HER2 therapy, trastuzumab (Wang *et al*., 2019). Our single cell RNA-seq data reveal downregulation of IFITM3 within 24 hours of HER2 overexpression, that is maintained until at least 72 hours, which could show that this does not decrease as a result of resistance but may predispose resistance to therapies at the very early stage of disease. Overall, our data show the power of combining genome-wide molecular approaches using an *in vitro* transformation model system to uncover subtle but relevant variations in cellular states. Given the dramatic remodelling of the chromatin state driven by a single factor in HER2 positive breast cancer, we speculate that other cancer types may also feature similar mechanisms of cellular transformation through chromatin remodelling. Cataloguing early chromatin changes can emerge as a promising therapeutic target, with a particular focus on early and low HER2 induced alterations in breast cancer.

Metastasis is multi-step, low probability process, in which primary cells must invade the local tissue and extravasate into a distant site. Our work shows that low HER2 expressing cells gain transformational ability through de-differentiation and dramatic chromatin remodelling. This model could be further extended to assess how low HER2-driven changes in chromatin state are used as a route for metastasis in *in vivo* models, and if low loss of differentiation correlates with aggressiveness in more physiologically relevant models.

## Materials and Methods

### Cell culture

The immortalised human mammary epithelial cell line MCF10A was obtained from the American Type Culture Collection (ATCC) and grown under recommended conditions. Briefly, MCF10A cell medium consists of Dulbecco’s Modified Eagle’s Medium (DMEM/F12) (SIGMA #D8347) supplemented with 5% Horse Serum (SIGMA #H1138), 0.5 μg/mL Hydrocortisone (SIGMA #H0888), 20 ng/mL Epidermal Growth Factor (EGF) (SIGMA #E4127), 100 ng/mL Cholera Toxin (SIGMA #C8052), 10 μg/mL Insulin (SIGMA #i9278) and 1X Pen/Strep.

HEK293T cells were cultured in Dulbecco’s Modified Eagle’s Medium (DMEM) (SIGMA#D5796) in 10% foetal bovine serum (FBS) with 1X Pen/Strep.

For 3D overlay cell cultures, cells were grown in chamber wells in a mixture of matrigel (CORNING #356230) and collagen (CORNING #11563550), which were mixed with 0.1M NaOH and 10X PBS, as previously described (Xiang and Muthuswamy, 2006). To collect cells from 3D cell cultures, cell recovery solution (CORNING #354253) was used at 4°C for 30 to 60 minutes according to the manufacturer’s instructions. Staining 3D acini were fixed with 4% paraformaldehyde (PFA). Acini were permeabilised with 0.5% Triton-X and blocked in 10% goat serum in PBS-Tween. Acini were stained with Phalloidin dye overnight at 4°C. The detachable chambers were removed, and acini mounted in mounting media reagent and allowed to dry in the dark at room temperature. Once dried, slides were visualised using a fluorescence microscope.

### Vectors and Viral infections

To generate HER2 inducible MCF10A cell line (Carter *et al*., 2017), we first transiently transfected HEK293T cells using jetPRIME transfection reagent (POLYPLUS #114-15). The inducible HER2 construct (ADDGENE #46948) alongside pMD2.G (ADDGENE #12259) [envelope plasmid], and of pCMV delta R8.2 (ADDGENE #12263) [packaging plasmid] were transfected into 90% confluent HEK293T cells for 24 hours. Lentiviral particles were harvested by centrifugation and early passage MCF10A cells were infected for 48 hours. Cells were then flow sorted based on GFP expression to obtain a pure population.

### Western blotting

Cells were harvested and lysed in RIPA buffer containing protease and phosphatase inhibitors. Lysates were mixed with sample loading buffer and proteins were resolved on sodium dodecyl sulphate-polyacrylamide gel electrophoresis (SDS-PAGE) and transferred onto PVDF membranes. Membranes were blocked in 5% milk, and antibodies were incubated overnight in 5% BSA solution. Antibodies used include HER2 (CELLSIGNALLING #2165), GAPDH (CELLSIGNALLING #2118), p53 (CELLSIGNALLING #2527), p27 (CELLSIGNALLING #3836), p21 (CELLSIGNALLING #2947), Tubulin (ABCAM #7291), Anti-rabbit secondary (Amersham ECL Rabbit IgG, HRP-linked whole Ab #NA934).

Human samples were obtained from Barts Cancer Institute tissue bank, where human samples were used, informed consent was obtained from all individual participants included in the study.

### Flow cytometry and flow sorting

Cultured cells were detached from plates with trypsin and stained with 2% horse serum. Cells were then stained with the following conjugated antibodies: HER2 (BD BIOSCIECNES #745299, 1:100), EpCAM (BD BIOSCIENCES #347200, 1:40), MUC1 (BD BIOSCIENCES #743410, 1:50), CD24 (BIOLEGEND #311135, 1:50) for 20 minutes at room temperature. Cells were washed in 1 mL of 2% horse serum and then resuspended in DAPI buffer. Stained cells were analysed on LSR Fortessa. For cell sorting, cells were stained with the antibodies of interest and isolated using ARIA fusion cell sorter.

### ATAC-seq library preparation and differential analysis

5×10^**5**^ cells were directly recovered from cell culture by trypsin from 2D cell culture or by using the recovery solution (CORNING #354253) for cells grown in either 2D or 3D cell culture. ATAC-seq libraries were generated as described previously (Buenrostro *et al*., 2015), with minor amendments. We performed 10 initial PCR amplification cycles followed by direct purification of the transposed DNA, without performing qPCR to calculate the additional numbers of required cycles. Sequencing data was aligned to the human genome (grch38) using bowtie2. Peaks were called on each biological replicate of all ATAC-seq reads using MACS2, and putative copy number and mitochondrial regions were removed. Peak dataset for differential analysis was generated by applying a threshold using a desired fold-change and a −log10 transformed FDR adjusted *p*-value. Differential accessibility was assessed using DiffBind and regions were called differentially accessible based on log2Fold change and FDR p-value.

### Phosphoproteomic sample preparation

For phosphoproteomic experiments, cells were grown in 2D cell cultures. Cell pellets were lysed using 8M urea lysis buffer (containing phosphatase inhibitors). The amount of protein in the lysates was quantified by BCA assay. 250 μg from each sample were digested into peptides with immobilised TPCK-trypsin beads (Thermo Fisher Scientific #20230) at 37°C overnight. Phosphorylated peptides were enriched from total peptides using TiO2 chromatography, as reported previously (Montoya *et al*., 2011; Larsen *et al*., 2005). Finally, peptides were snap frozen and dried in a SpeedVac. Dried peptides were dissolved in 0.1% TFA and analysed by LC-MS/MS on Q Exactive plus mass spectrometer (Thermo Fisher Scientific). Peptide identification was performed using the Mascot search engine (Casado and Cutillas, 2011). Allowed variable modifications were phosphorylation on Ser, Thr and Tyr, and oxidation of Met, and the Pescal software (Casado and Cutillas, 2011) and (Cutillas and Vanhaesebroeck, 2007) was used to quantify the peptides. Kinase-substrate enrichment analysis (KSEA) (Casado *et al*., 2013) was used to determine kinase activities. The intensity values were calculated by determining the peak of each individual extracted ion chromatograms (XIC) and plotted as heatmaps. The resulting quantitative data were transferred and visualised in Microsoft Excel. The significance (log2 fold change < −0.5-fold, FDR corrected p-value of < 0.05 for downregulated phosphosites and log2 fold change > 0.5-fold, FDR corrected p-value of < 0.05 for the upregulated phosphosites) of each phosphosite was annotated by an asterisk; we used the “filter” function on excel to filter out those phosphosites that were not significant. All of the significant MCF10A^GFP^ data was filtered out, whilst simultaneously filtering out non-significant data for the MCF10A^HER2^ cells, giving us significant changes in MCF10A^HER2^ cells that are not significantly changing in the MCF10A^GFP^ cells. The number of phosphosites was determined by the number of columns as each column contains one phosphosite, unless overlapping sites were present, in which case they were manually counted.

### Migration/Invasion assays

Chilled matrigel or collagen mixture was directly pipetted on the centre of 8 μm pore size transwell inserts (MILLICELL #MCEP12H48) that were placed into a 12-well plate, and allowed to solidify at 37°C. Meanwhile, cells were trypsinised and pipetted onto the transwell inserts – which were either coated with matrix or left uncoated – and cultured for 16 hours. Highly migratory/invasive cells were stained with 0.05% of crystal violet dye. Images of random regions were taken using a standard light microscope and quantified using imageJ.

### Soft agar colony formation assays

A 0.8% base layer was formed in plates using ultra-pure culture grade agarose (THERMOFISHER #16500500) allowed to settle at room temperature. 5000 cells per well were mixed with 0.3% agarose and plated evenly, drop-wise, on top of the base layer. Media was changed every 2 days for 3 weeks. Colonies were fixed using 4% paraformaldehyde (PFA) and permeabilised using 100% methanol. Colonies were stained using 0.05% crystal violet dye and images were taken using a dissecting microscope. Binary masks were applied to each of the images and thresholding parameters for the diameter ranging from 10um 100μm were set on ImageJ. Colonies were counted using ImageJ only if they satisfied criteria above the threshold values, and colony counts were then manually checked and adjusted if necessary.

### Single cell RNA seq

MCF10A cells were induced with 1μg/ml of doxycycline 0, 7, 24, 48 and 72 hours in 2D cell cultures. Cells were then detached using TrypLE (Gibco) and collected in 1X DPBS (Gibco). After one wash in 1X DPBS, cells were resuspended in 2% BSA-DPBS at a concentration of 10,000 cells/μl. 500 000 cells (50μl) were blocked with 10μl TruStain FcX blocking solution (BioLegend). Each treatment group was stained with 0.5μl of specific Totalseq-A Hashtag antibodies and 0.5μl of TotalSeq™-A0133 anti-human CD340 (ERBB2/HER2) protein expression antibody. Cells were washed 3 times with 1 ml 2% BSA-DPBS and resuspended to a concentration of ~10,000 cells/μl. Equal volumes of each treatment group were pooled, and cell pool was assessed for cell concentration and viability. Single-cell cDNA, protein expression (ADT) and hashtag (HTO) libraries were generated using Chromium Single Cell 3’ version 2 reagents (10x genomics and Biolegend) per manufacturers’ protocols. Single-end sequencing of libraries was performed by Novogene Corporation Inc. on a Novaseq 6000 (Illumina) sequencer with HTO libraries constituting 5% of the sample.

Single cell data was run through the 10X cellRanger pipeline to produce counts tables for gene expression counts, HTO counts for sample identification and ADT counts for HER2 expression. Cells were identified and assigned to a timepoint using the HTO counts table and the HTODemux method in Seurat. To exclude cells that did not respond to the doxycycline induction, treated cells with less than 35 counts of the ERBB2/HER2 expression tag were filtered out. The remaining gene expression data was run through Seurat’s basic data processing pipeline. The data was normalized, scaled, and the effects of cell cycle were regressed out using Seurat’s cell cycle regression strategy. The data was then run through principal component analysis (PCA). The principal components were used to identify clusters and UMAP was run for visualization. Two different differential expression analyses were run using Seurat’s FindAllMarkers function, one across the different clusters and one across the different timepoints.

## Supplementary material

**Supplementary Figure 1.**
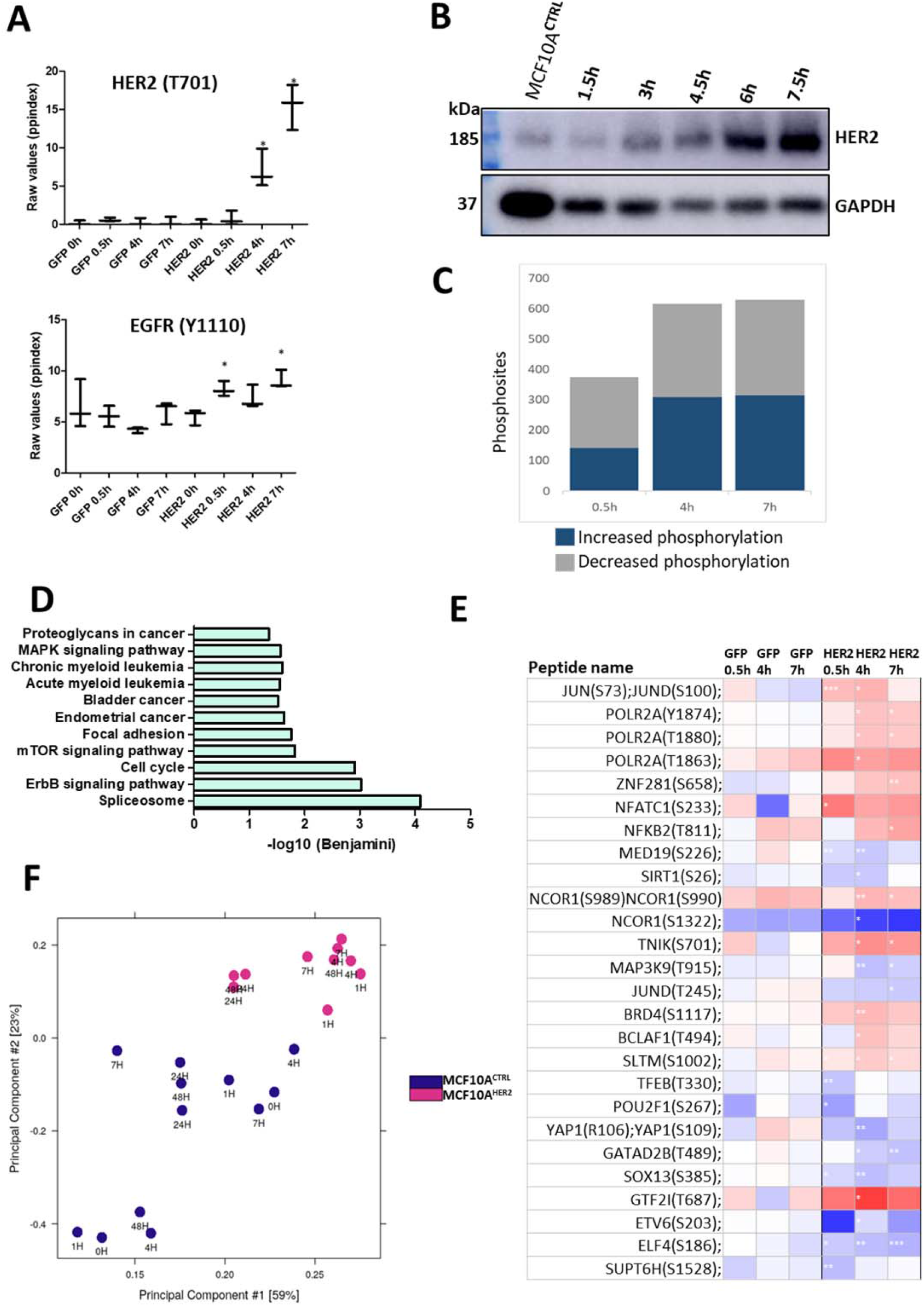
(A) An internal quality control (QC) for phosphoproteomic analysis. HER2 phosphorylation modification (T701) increases in a time dependent manner. EGFR [HER1] (Y1110), a family member of HER2, also becomes marginally activated in a time dependent manner compared to control cells. [* FDR corrected p-value of < 0.05]. (B) Western blot analysis of HER2 protein in a time-dependent manner in the early time-points upon induction with the same concentration of doxycycline (1μg/ml) from 0h to 7.5h. GAPDH was used as a loading control. N=1. (C) Bar graph depicting the number of detected phosphosites increasing or decreasing in phosphorylation in the phosphoproteomic analysis at the time-points analysed. Significance is shown to log2fold change > 0.5, FDR corrected p-value of < 0.05. This graph shows analysis performed using lower statistical threshold compared to figure 2B. (D) Signalling pathway analysis of the early immediate changes in transformation. Signalling pathway analysis using the DAVID Bioinformatics tool of the differentially phosphorylated events at all time points investigated upon HER2 protein induction is shown. Significance threshold applied here; log2fold change > 0.5, FDR corrected p-value of < 0.05. (E) Identification of transcription factors and chromatin regulators. A list of transcription factors and chromatin regulators becoming differentially phosphorylated upon HER2 expression in at least one time-point upon HER2 overexpression but are not significantly changing in GFP-transduced MCF10A cells. [* FDR corrected p-value of < 0.05, **FDR corrected p-value of < 0.001, *** FDR corrected p-value of < 0.001]. (F) Principal component analysis (PCA) of all samples used in this study. Samples are colour-coded by cell type.

**Supplementary Figure 2.**
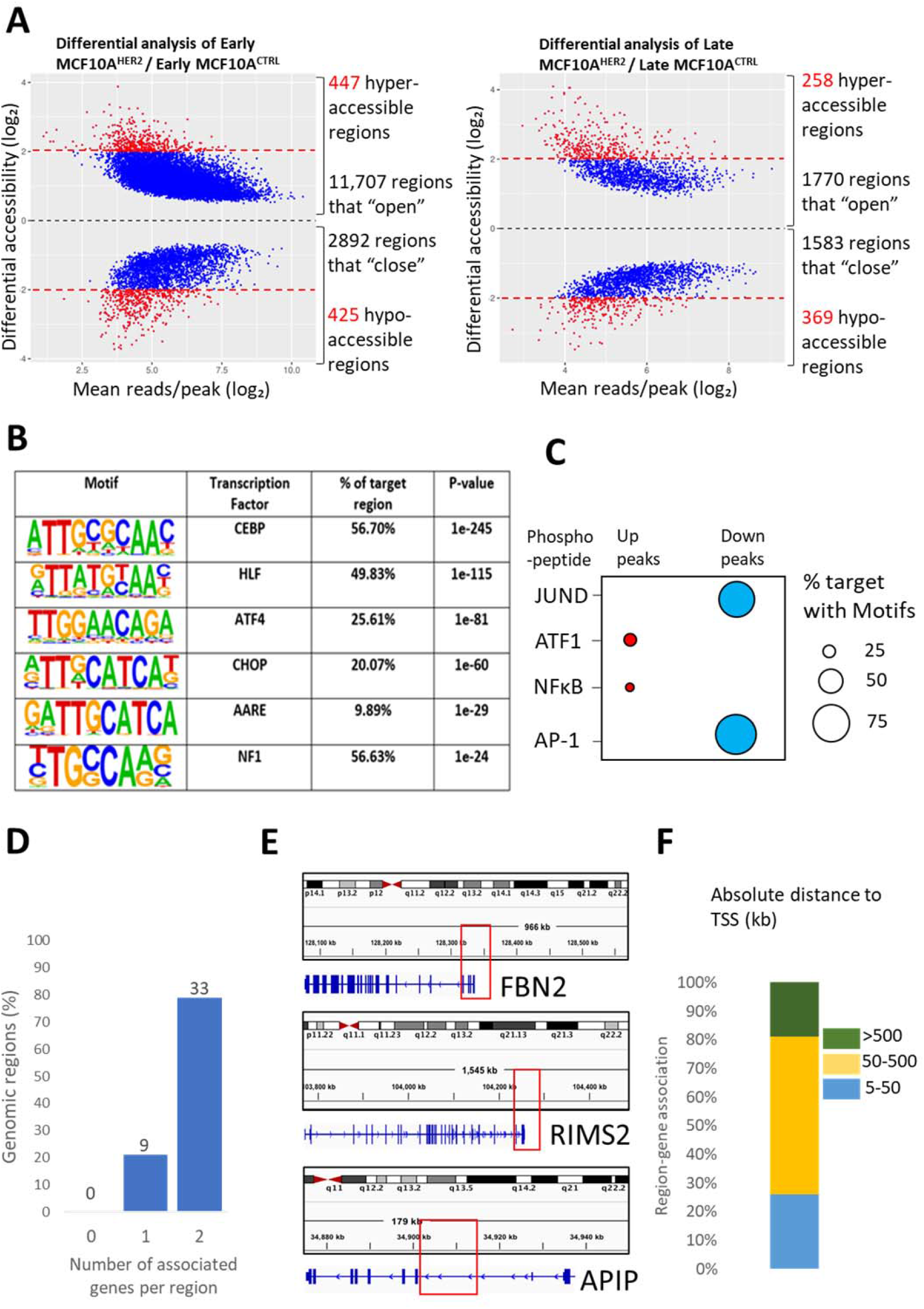
(A) Differential accessibility (log2fold change > 0.5, FDR corrected p-value of < 0.05) shown by MA plot between MCF10A^**HER2**^ and control cells, plotted against the mean reads per region. Cells were grown in 3D cell culture from 0-48 hours and ATAC-seq performed on their acini. “Early” time-points represents 0h, 1h, 4h, and 7h data combined. “Late” time point represents 24h and 48h time-points combined. Each dot represents a region, with the blue dots representing a log2fold change of at least 0.5. (B) Enrichment of transcription factor recognition sequences in differential ATAC-seq peaks comparing MCF10A^HER2^ and control cells based on HOMER analysis using the accessible (up) peaks. (C) Motif analysis from ATAC-seq reveal shared transcription factors identified in the phosphoproteomic screen. A dot plot of the overrepresented motifs in differentially accessible regions with the size of the circle representing the % of differentially accessible regions that contain the motif in the accessible and inaccessible peaks. (D) GREAT database analysis showing the number/percentage of genes associated per region of the common regions found between the early down and late down peaks in the ATAC-seq data. (E) Integrative Genome Browser (IGV) panel displaying regions associated with the indicated genes. Red boxes show chromosomal coordinates (128,340,997-128,341,448) found in ATAC-seq data, which associates with the promoter region of FBN2. (F) Absolute distance to closest transcription start sites (TSSs) of the common differentially inaccessible regions.

**Supplementary figure 3.**
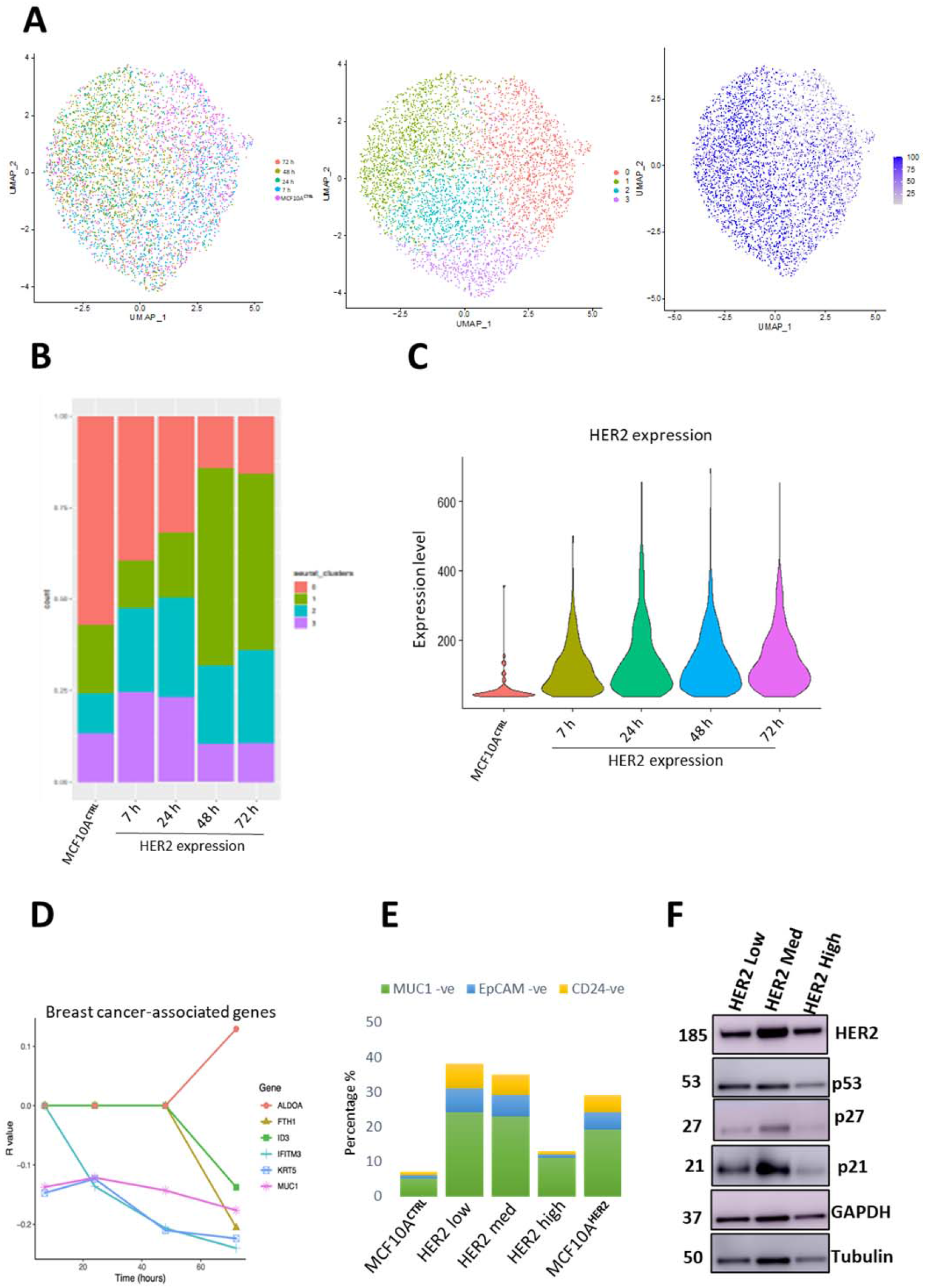
(A) UMAP plot showing clustering based on different time points. UMAP plot displaying clusters of genes with similar features. UMAP plot showing a range of HER2 gene expression. (B) Bar graph showing Seurat clustering which defines clustering via differential gene expression. (C) Violin plot shows HER2 levels increase in a time-dependent manner with HER2 expression. (D) Single cell RNA sequencing was performed on MCF10A cells with HER2 induction from 0 to 72 hours (3 days). Line graph shows R values as a measure of linear relationship between HER2 expression increase (with time) and some genes of interest that either increase in expression or decrease in with HER2 expression (E) Cells were analysed by flow cytometry and HER2 positive cells were separated into three subpopulations of low, medium, and high HER2 overexpression as indicated. The enrichment of stem markers is shown as a proportion of the total number of cells exhibiting MUC1 ^−ve^, EpCAM ^−ve^ and CD24 ^−ve^ phenotype. (F) Western blot of the indicated proteins known to have higher expression in cells that have undergone OIS. Protein lysates were prepared from cells sorted based on HER2 expression. HER2 was induced in cells for 3 days (MCF10A^HER2^) and then FACS separated based on HER2 expression into three different subtypes (low, medium, and high HER2 expressing cells). GAPDH and Tubulin were used as loading controls. N=3.

**Supplementary figure 4.**
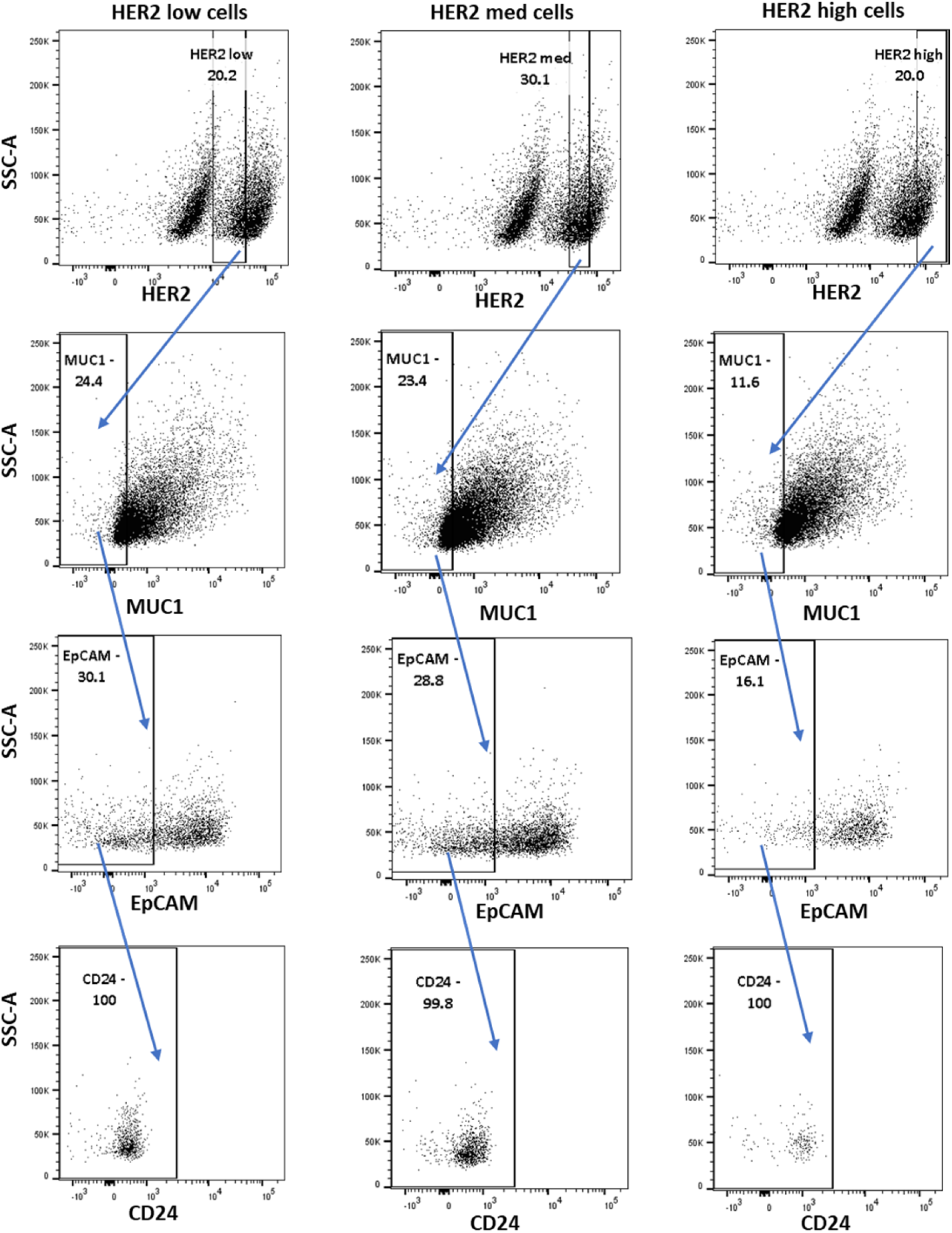
Cells were analysed by flow cytometry and HER2 positive cells were separated into three subpopulations of low, medium, and high HER2 overexpression as indicated. The enrichment of stem markers is shown as a proportion of the total number of cells exhibiting MUC1 -ve and EpCAM -ve and CD24 -ve phenotype. The blue arrows indicate step-by-step analysis of the HER2 subpopulations, and the respective enrichment of breast stem markers in each subtype.

### ATAC-seq bioinformatics analysis pipeline

The ATAC-seq data was provided as FASTQ files. Quality control of raw sequencing read files was performed using FastQC. Illumina adapter trimming was done using Cutadapt; settings: Cutadapt -a CTGTCTCTTATACACATCT -A CTGTCTCTTATACACATCT -o out.1.fastq -p out.2.fastq. Trimmed reads were aligned using the human genome, Genome Reference Consortium Human Build 38 patch release 13 (GRCh38.p13), using bowtie2, and a SAM file was obtained; setting: bowtie2 index −1 trimmed FASTQ file −2 trimmed FASTQ file –S 1.sam. The resulting sam files were converted into binary bam files; setting: Samtools view –Sb in.samfile > out.bamfile and sorted; setting: Samtools sort in.bamfile -o out.bamfile and indexed; setting: Samtools index in.bamfile. To ensure an improved mapping quality, we removed mitochondrial DNA; setting: Samtools view –h in.bamfile | removeChrom - - chrM | Samtools view - b - > out.bamfile. PCR duplicates were removed from the files using Picard tools; setting: Java -jar picard.jar MarkDuplicates I=in.bamfile O= out.bamfile M=dups.txt REMOVE_DUPLICATES=true VALIDATION_STRIGENCY=LENIENT.

For viewing samples on genome bowser or assessing reproducibility and data exploration, all samples were “down sampled” to the same number of reads; setting: samtools view -b -s [downsampling_ratio] in.bam > out.downsampled.bam. Peaks calling was done for each individual non-downsampled file with MACS2 “callpeak”; setting: MACS2 callpeak -t inbamfile -f BAMPE -n in.bamfile -g ce –keep-dup all. These files were then analysed using DiffBind for differential analysis on R. For each sample, a path to the peaks and the bam file were listed in Microsoft Excel and loaded in R; setting: db.object = dba(sampleSheet=“name_of_sample_sheet”). Then, the next step was to take the alignment files and compute count information for each of the peaks/regions in the consensus set; setting: db.object = dba.contrast(db.object, categories=DBA_TREATMENT, block=DBA_CONDITION, minMembers = 2); setting: db.object = dba.analyze(db.object,bParallel=TRUE,method=DBA_ALL_METHODS). R was used to plot the differential changes such as MA plot with an appropriate threshold; setting: dba.plotMA(db.object,th=“0.05”,method=DBA_DESEQ2). Significant changes could then be saved from up or down peaks e.g.; setting: up_peaks_db.object.SigChanges.0.05FDR <- db.object.SigChanges.”0.05FDR”[db.object.SigChanges.0.05FDR$Fold > 0,] and counted using the command line and can be plotted as percentage in Prism or Microsoft excel in the form of a chart/graph.

## Acknowledgements

We thank Dr Salvatore Federico Pedicona and Dr Hemalvi Patani for critically reading the manuscript. We thank Kriszta Kovacs for her assistance in 3D morphology assays. This work was supported by a Leverhulme Trust grant.

